# CryoPhold: CryoEM meets AlphaFold and molecular simulation to reveal protein dynamics

**DOI:** 10.1101/2025.09.12.675912

**Authors:** Soumendranath Bhakat, Shray Vats, Andreas Mardt, Eva M. Strauch

## Abstract

Here we are introducing CryoPhold, a modular workflow that unifies AlphaFold-based ensemble generation, Bayesian reweighting against experimental cryo-EM maps, molecular simulation, and machine learning to quantify conformational populations and identify structural fingerprints that govern protein functions. Proteins are inherently dynamic, interconverting among conformational states that govern their function. Perturbations such as mutations, ligand binding, and pH changes modulate these dynamics and are implicated in many diseases. While cryogenic electron microscopy (cryo-EM) has transformed structure determination, it typically yields an averaged density map representing a static snapshot. A central challenge remains capturing the thermodynamics underlying protein motions and corresponding structural fingerprints that modulate function. CryoPhold enables Bayesian reweighting of AlphaFold-generated structural ensembles against experimental cryo-EM maps, generating posterior structural ensembles that are consistent with experimental data while preserving conformational heterogeneity. Molecular simulations seeded from the posterior ensemble capture time-dependent dynamics, while machine learning models trained on featurized molecular simulation data identify structural fingerprints (“hotspots”) that modulate protein dynamics. Finally, Markov state models trained on featurized molecular simulation data quantify metastable state populations and free-energy landscapes. By integrating a generative AI-based protein structure prediction model, experimental cryo-EM density, physics-based sampling, and machine learning, CryoPhold enables dynamics paradigm to capture biomolecular motion. The workflow enables prediction of equilibrium populations and structural fingerprints governing conformational dynamics in human transporter protein, GlyT1. It further captures structural changes and population shifts associated with oncogenic BRAF mutants, key driving factors behind melanoma progression.

## Introduction

Proteins are inherently dynamic entities whose biological functions depend on their ability to interconvert among multiple conformational states spanning a vast range of timescales—from sub-nanoseconds to seconds^1^. These motions modulate essential processes such as ligand recognition, allosteric regulation, and catalysis. Precisely characterizing this conformational heterogeneity, however, remains one of the foremost challenges in structural biology^2^.

Cryogenic electron microscopy (cryo-EM) has revolutionized macromolecular structure determination, consistently providing near-atomic resolution density maps for stable protein-protein complexes. However, a significant challenge arises with *apo* proteins (proteins not stabilized by interactions with other proteins, RNA/DNA, ligands or other environmental stimuli). For these, even advanced cryo-EM reconstructions often result in blurred, low-resolution, or noisy densities in highly dynamic regions. These regions are frequently where motions critical for biological function occur. Further, the flexibility of apo proteins presents a bottleneck when fitting their structures into cryo-EM maps as X-ray crystal structures of *apo* proteins often lack domains that are highly dynamic^3^.

Current computational approaches attempt to reconcile structural models with cryo-EM maps through mainly three different categories: a) rigid-body^4^ or flexible^5^ fitting of available X-ray structures, b) normal-mode^6,7^ or neural network-based schemes^8,9^ that capture *fast* internal motions from heterogeneous single-particle cryo-EM datasets, and c) density-restrained molecular-dynamics simulations^10,11^ that refine coordinates against the map. All three strategies fail when no suitable starting structure exists or when the region of interest is highly dynamic and poorly resolved. Additionally, fast domain motions recovered from dimensionality reduction techniques such as normal-mode analysis or principal component analysis (PCA) do not capture slowly varying conformational motions in biomolecules, which are directly connected to interchanging metastable states governing protein ^12^ functions^13^.

Recent studies demonstrate that lowering the depth of the multiple-sequence alignment fed to AlphaFold2 (AF2)^14^ yields a heterogeneous conformational ensemble^15–17^. Whether this ensemble encompasses physical information, statistical patterns in the training data, or outright memorization remains an open question^18–20^. However, such a conformational ensemble provides a starting hypothesis of the dynamical region that remains unresolved by X-ray crystallography. The key challenge is to merge the AF2 generated ensemble with experimental cryo-EM data and physics-based simulations, then extract the *slowly* varying conformational features that govern protein dynamics^21,22^.

Introducing CryoPhold, a first-of-its-kind modular workflow that enables conformational ensemble generation by seamlessly integrating AF2 predictions with reference cryo-EM density maps, while simultaneously predicting conformational populations and identifying structural hotspots by integrating cryo-EM refined conformational ensembles with molecular simulations and machine learning.

The modular workflow consists of the following steps: **(1)** MSA subsampling combined with AF2 produces an initial structural ensemble of apo protein from its sequence, **(2)** Bayesian reweighting against the cryo-EM density map retains structural models consistent with the experimental data while preserving conformational diversity. **(3)** physics-based molecular simulations^23^, seeded from this reweighted posterior ensemble, captures temporal evolution of dihedral angles and distances which encodes conformational dynamics. **(4)** a ML-module consists of slow feature analysis (SFA)^24,25^ and a time-lagged autoencoder (TAE)^22,23^ discovers the slow, functionally relevant motions and corresponding structural features, mapping them to a latent space (**Figure 1**). Markov state model (MSM)^26–29^ trained on the key structural features quantifies state populations, transition times, and the underlying free-energy landscape.

**Figure 1.**
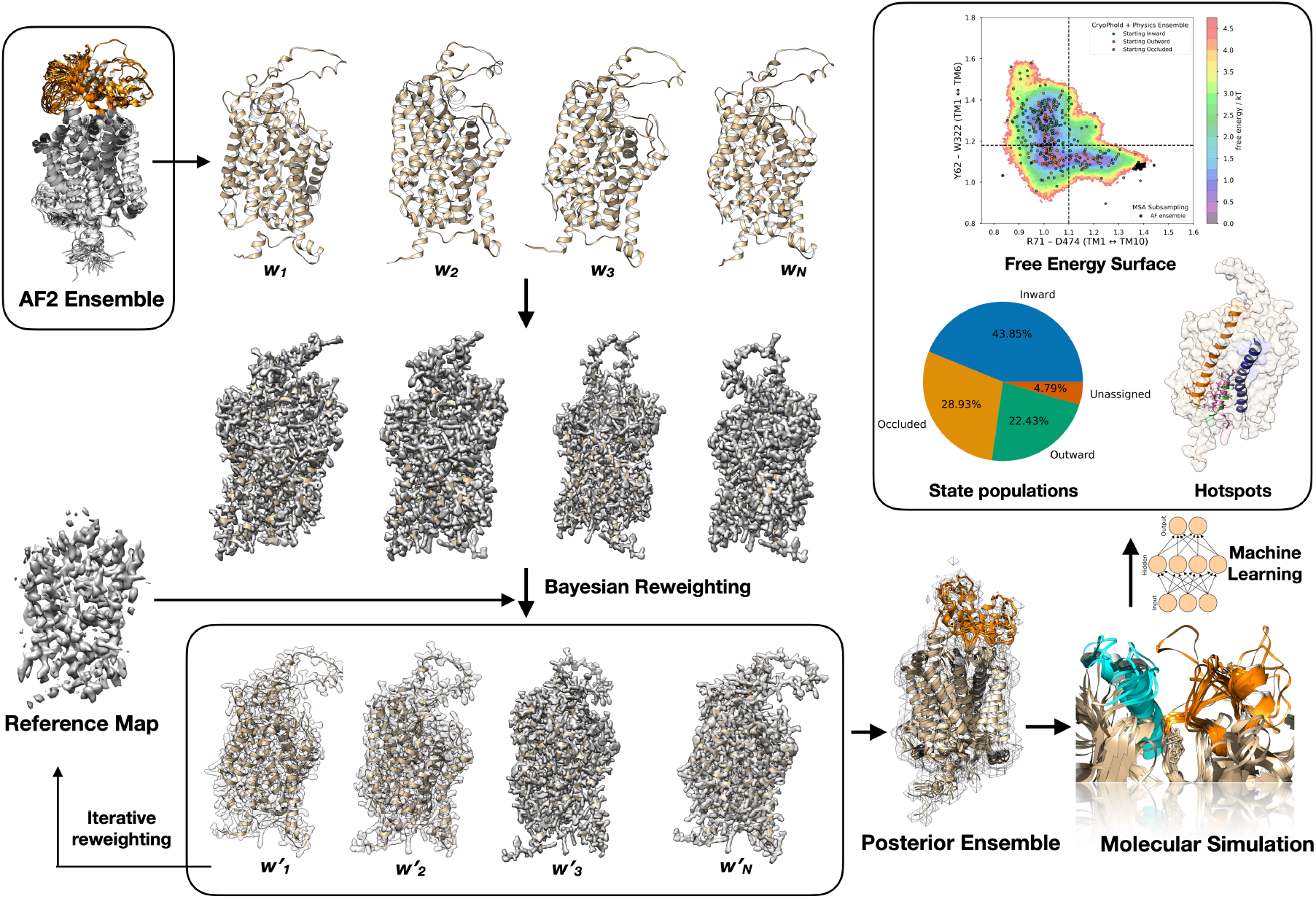
Schematic illustration of the CryoPhold workflow. The prior ensemble, generated by MSA subsampling of AF2, is aligned to the experimental cryo-EM map. Bayesian reweighting assigns posterior weights (w′_i_), yielding a “best fit” and posterior ensemble (highlighted by different shades of map post reweighting) consistent with the reference map. Molecular simulations initiated from this ensemble and analyzed with Markov state models (MSMs) capture equilibrium populations of metastable states (pie chart) and free energy surface along key structural features that govern biomolecular dynamics. Machine learning trained on the simulation data identifies conformational hotspots and *slowly fluctuating* structural features underlying conformational transitions.

By combining AF2, cryo-EM density, physics-based sampling, and machine learning, our workflow captures biomolecular dynamics that extend far beyond static structural snapshots, enabling the era of ensemble paradigm for downstream applications such as ensemble-based virtual screening or dynamics-aware therapeutics design^30^.

## Results

Identifying conformational ensembles and populations of different states in *apo* proteins, which can adopt multiple conformations in response to ligand binding, as well as predicting population shifts upon mutations, are two key challenges in structural biology. We chose membrane transporter, Glycine transporter 1 as a model system for the first case, for which holo states and corresponding cryo-EM structures have been resolved. Monomeric kinase, BRAF and its oncogenic mutations (V600E and V600K) were chosen for the second case, where cryo-EM structures of the corresponding proteins have been resolved but lacks an indication of population shifts upon mutations.

### CryoPhold captures conformational dynamics in glycine transporter 1 (GlyT1)

Glycine transporter 1 (GlyT1)^14^, a member transporter of the solute carrier 6 (SLC6) family^31^, modulates neurotransmission by clearing glycine from the synaptic cleft. Through glycine level control, GlyT1 indirectly regulates the activity of the N-methyl-D-aspartate (NMDA) receptor. Growing evidence links NMDA receptors to schizophrenia development, making GlyT1 an attractive target for novel schizophrenia treatments^14,31^.

Cryo-EM and X-ray crystallography studies have revealed three distinct GlyT1 states: (a) occluded (PDB 8WFI; EMDB: EMD-37492; 2.58 Å), bound to glycine; (b) inward (PDB 8WFJ; EMDB EMD-37493; 3.35 Å), bound to sarcosine-based inhibitors like ALX-5407; and (c) outward, bound to non-sarcosine inhibitors such as SSR-504734 (PDB 8WFK; EMDB: EMD-37494; 3.22 Å) and PF-03463275^14^.

However, understanding the conformational equilibrium among metastable states in apo GlyT1—and identifying the structural hotspots that control these dynamics—remains difficult. Additionally, extracellular loops in GlyT1 are only partially resolved in static structures because of their inherent flexibility. Therefore, we investigated whether we could infer populations and key structural features influencing apo GlyT1 dynamics using an AF-generated conformational ensemble and cryo-EM maps as priors.

A prior ensemble was generated by MSA subsampling integrated with AF, yielding 80 structurally diverse models. Applying a pLDDT based filtering (see “Filtering” subsection in Methods) removed erroneous predictions and retained 50 models while preserving conformational heterogeneity. Each model was rigid-body aligned to cryo-EM maps representing occluded, inward, and outward states using the standalone Situs package, and the aligned structures with their corresponding maps were incorporated as priors within the CryoPhold module. This yielded a posterior ensemble of 34 structures (occluded: 8; inward: 10; outward: 16). Independent molecular dynamics simulations (100 ns each) were initiated from every CryoPhold derived structure, generating a combined dataset of 3.4 µs. The aggregated trajectories were used to construct a MSM (see Methods). Equilibrium populations from the MSM, projected onto Cα–Cα distances Y62 (TM1)–W322 (TM6) and R71 (TM1)–D474 (TM10), captured the collective motions of TM1, TM6, and TM10 underlying transitions among occluded, inward, and outward states. CryoPhold-augmented molecular simulations revealed that apo GlyT1 samples a dynamic equilibrium across multiple metastable states, with populations ordered as *inward > occluded > outward* (**Figure 2**).

**Figure 2.**
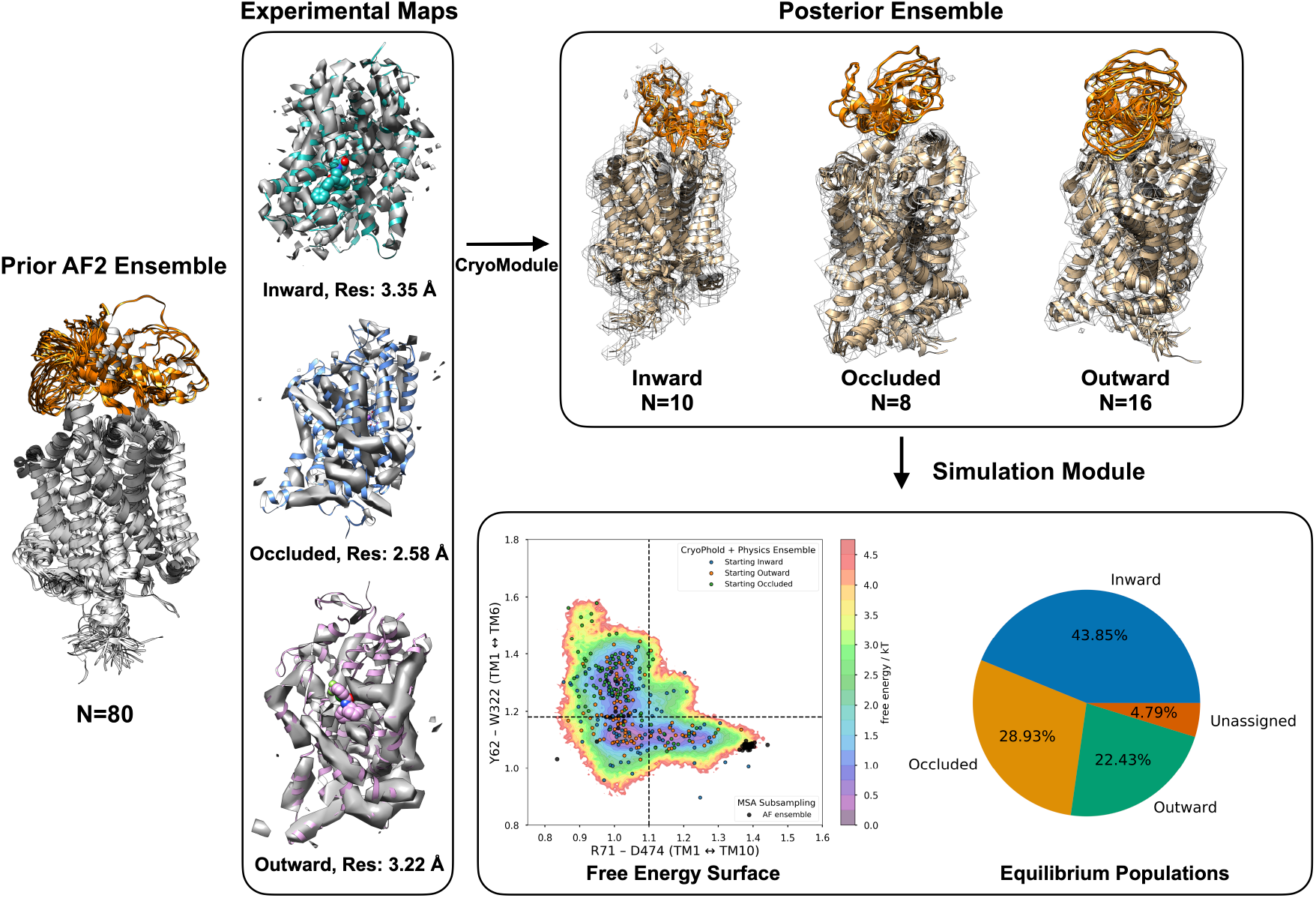
Overview of the CryoPhold workflow applied to apo GlyT1. Cryo-EM structures of GlyT1 have been resolved in three states: inward (ALX-5407 bound), occluded (glycine bound), and outward (SSR-504734 bound),. The small molecules, ALX-5407 and SSR-504734 are represented as spheres. Bayesian reweighting of reference cryo-EM maps with a prior ensemble generated by MSA subsampling of AF2 yields smaller posterior ensembles that best match these states. The simulation module enables molecular dynamics simulations from the posterior ensemble, which samples dynamics along low-dimensional collective variables i.e Cα–Cα distances between Y62 (TM1)–W322 (TM6) and R71 (TM1)–D474 (TM10). By contrast, MSA subsampling alone (black dots) failed to capture conformational transitions or metastable states in apo GlyT1. The extracellular loop of GlyT1 is highlighted in orange which is not resolved in experimental cryo-EM and X-ray crystal structures.

A time-lagged autoencoder (TAE)^13^ trained on χ1 and χ2 dihedral angles from the MD data generated by the simulation module, identified slowly varying coordinates and highlighted residues critical for apo *GlyT1* dynamics (**Figure 3**). Among these, it identified Y62, a residue that functions as a “*lid*,” whose flipping is required for binding of the sarcosine-based inhibitor ALX-5407 and drives a shift from the occluded to the inward state. Consistent with this prediction, *Wei et al*. reported that the Y62A mutation reduces glycine uptake activity, underscoring the functional role of Y62 in GlyT1^14^. Furthermore, SFA was trained on Cα– Cα distance pairs, identified slowly varying contacts governing apo GlyT1 dynamics, and, from first principles, recovered binding hotspots for both sarcosine-and non–sarcosine–based inhibitors.

**Figure 3.**
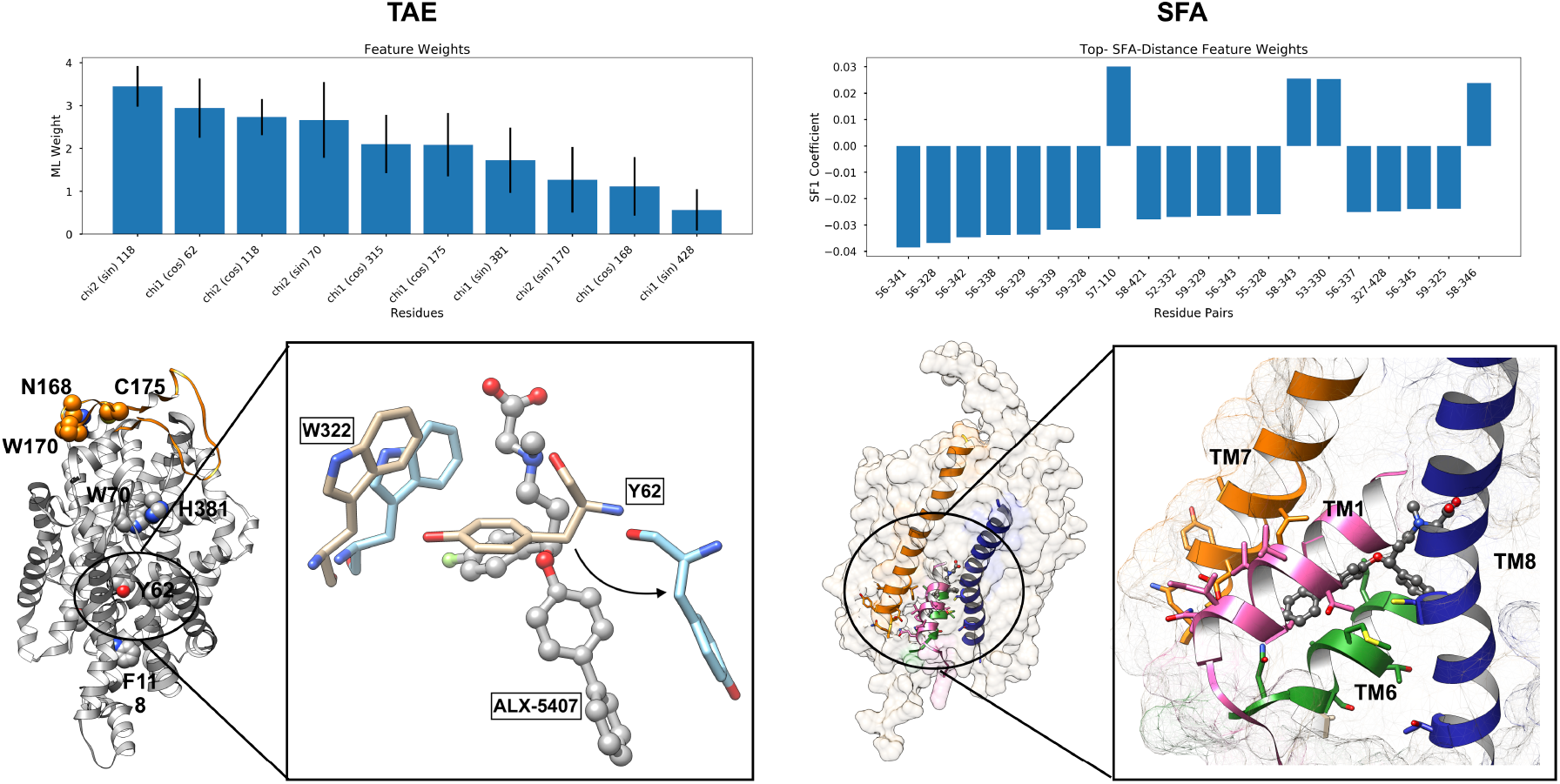
A time-lagged autoencoder (TAE) trained on χ1 and χ2 angles from CryoPhold-augmented molecular simulations identifies key residues whose flipping governs the dynamics of apo GlyT1. Among these, Y62 is critical: its flipping drives conformational changes that expose an allosteric pocket enabling binding of the sarcosine-based inhibitor ALX-5407. In parallel, the first component of slow feature analysis (SFA) trained on Cα–Cα distances highlights a conformational hotspot at the intersection of TM1, TM6, TM7, and TM8 that regulates transitions between metastable states and pocket opening. This integrated workflow is generalizable across a broad range of transporters.

It is important to note that, unlike small molecules stabilizing *outward* conformations by binding to exposed pockets in transporters, molecules that stabilize the inward conformation act through conformational allostery, as their binding sites are buried within the transmembrane helices. These allosteric transitions are driven by slowly varying structural rearrangements. The ability of CryoPhold to capture conformational hotspots governing such transitions in apo GlyT1 demonstrates the transferability of our workflow across different transporter protein families (Figure S2 in Supporting Information).

Together, this integrated pipeline quantifies state populations in apo GlyT1 and pinpoints structural hotspots that regulate its dynamics (**Figure 2**). We anticipate that this approach will extend to other transporters and membrane protein classes, providing a generalizable framework for uncovering conformational mechanisms across the transporter family.

### Wild-Type and oncogenic BRAF V600E/K exhibit divergent conformational dynamics

In normal cells, wild-type BRAF predominantly adopts a monomeric, autoinhibited state. In malignant melanoma, BRAF is mutated in ~66% of cases, with the single-point mutation V600E accounting for ~80% of these. V600E, located in the activation loop of monomeric BRAF, stabilizes the αC-in active conformation and renders BRAF constitutively active, continuously driving MAPK signaling and cancer cell proliferation. Understanding how oncogenic mutations modulate the conformational dynamics of monomeric BRAF is key to developing therapeutics against oncogenic BRAF variants^32–34^.

*Lavoie et al*. used cryo-EM to resolve monomeric BRAF WT and V600E in complex with 14-3-3, achieving maps at 3.29 Å (EMDB: EMD-43674) and 3.74 Å (EMDB: EMD-43673) resolution, respectively^33^.

However, the X-ray crystal structure of monomeric BRAF used to fit the V600E cryo-EM map (PDB: 8VYP) lacks large segments of the G-loop, β-strands, activation loop, and the active-site DFG loop. In the case of WT, the activation loop is also missing in the corresponding X-ray structure (PDB: 8VYO). These structural gaps, combined with the static nature of X-ray snapshots, limit a complete understanding of how the V600E activation-loop mutation drives a population shift (Figure 4).

**Figure 4.**
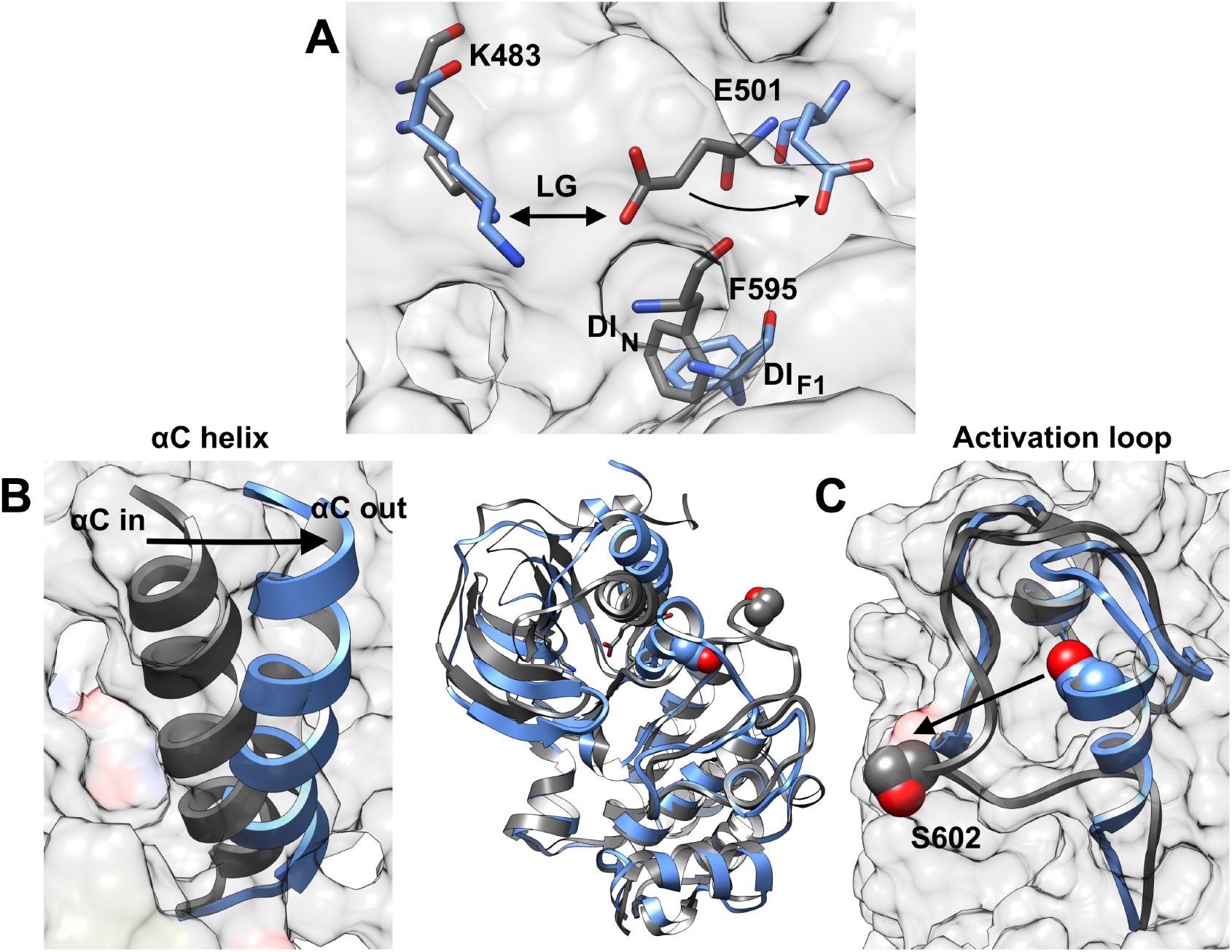
A: Relative positioning of DFG-Phe (F595) and the Lys–Glu (K483–E501) salt bridge in active (grey) and inactive (blue) conformations of monomeric BRAF. B: Active and inactive apo BRAF ensembles are distinguished by αC-helix in (active) and out (inactive) states, shown in ribbon representation (arrow highlights in to out transition). C: Activation loop transitions from folded (blue, AC_in_) to extended (grey, AC_out_) mark the shift from inactive to active states (arrow highlights transition from folded to the extended state). The Cα–Cα distance between S602 and D594 is used as a feature to quantify activation loop dynamics.

Conformational ensembles of monomeric WT and V600E were generated using the “*folding module*” by enabling MSA-subsampling with AF. UCSF Chimera^35^ was used to extract electron density corresponding to WT and V600E BRAF monomers from full cryo-EM maps, and rigid-body alignment of the AF2 structural ensemble for WT and V600E was performed against the focused electron density map of BRAF monomers (*see Methods for the alignment protocol*). The CryoPhold module was then applied to the aligned AF-derived ensembles to produce single best-fit models and posterior structural ensembles that maximally represents the cryo-EM maps. The posterior structural ensemble contains 11 and 16 PDB structures, respectively, for WT and V600E monomers.

The “*simulation module”* was used to launch 3 independent 200 ns MD simulations from each structure in the posterior ensemble, yielding a total of 6.6 µs for WT and 9.6 µs for V600E. Markov state models (MSMs) were constructed using the χ1 and χ2 dihedrals of DFG-Phe, and equilibrium populations were projected onto key structural features^23^ *(collective variables*) to capture the mutation-dependent population shift in monomeric BRAF.

We define the global DFG-in (**DI**) ensemble of BRAF by three states based on the DFG-Phe sidechain χ1 angle: DI_N_ (χ1 between −2 and 0 rad), DI_F1_ (χ1 between 0 and +2 rad), and DI_F2_ (χ1 < −2 rad or χ1 > +2 rad). Active serine/threonine kinases (STKs) such as BRAF predominantly occupy DI_N_, whereas inactive forms favor DIF_1_/DI_F2_. The *αC-helix in* → *out* transition is tracked by the *Lys–Glu* (**LG**) salt bridge: *latched* (LG_L_) when *αC* is “in” and *unlatched* (LG_U_) when *αC* is “out” (**Figure 4**). Each DI macrostate can comprise microstates in which the Lys–Glu interaction is either latched or unlatched (DI_N_LG_L_, DI_N_LG_U_, DI_F1_LG_L_, DI_F1_LG_U_, DI_F2_LG_L_, DI_F2_LG_U_) ^36^.

CryoPhold generated conformational ensemble captured the full BRAF monomer for both WT and V600E, resolving microstates defined by DFG-Phe flipping and *αC helix* in ↔ out transitions (**Figure 5**). Notably, the WT apo ensemble includes DI_F1_LG_U_ microstates, whereas V600E prefers DI_N_LG_L_—consistent with the αC-out and αC-in conformations observed in X-ray crystal structures. MSM analyses of the simulation data revealed that oncogenic V600E mutation shifts the DFG-Phe population from inactive (DI_F1_/DI_F2_) to active ensemble (DI_N_) and simultaneously captures αC-out (LG_U_) to αC-in (LG_L_) transition. This *inactive* → *active* shift in monomeric BRAF also altered activation-loop dynamics, with V600E mutation shifts the conformational equilibrium from folded (AC_in_) to extended (AC_out_) conformation (Figure 3C and Figure S1 in Supporting Information). TAE trained on the dihedral angles (ϕ, ψ, and χ1) extracted from the CryoPhold augmented molecular simulation identified F595 (DFG-Phe) and V/E600 as key residues governing BRAF conformational dynamics, along with several additional residues across the kinase N-lobe and C-lobe (**Figure 6**).

**Figure 5.**
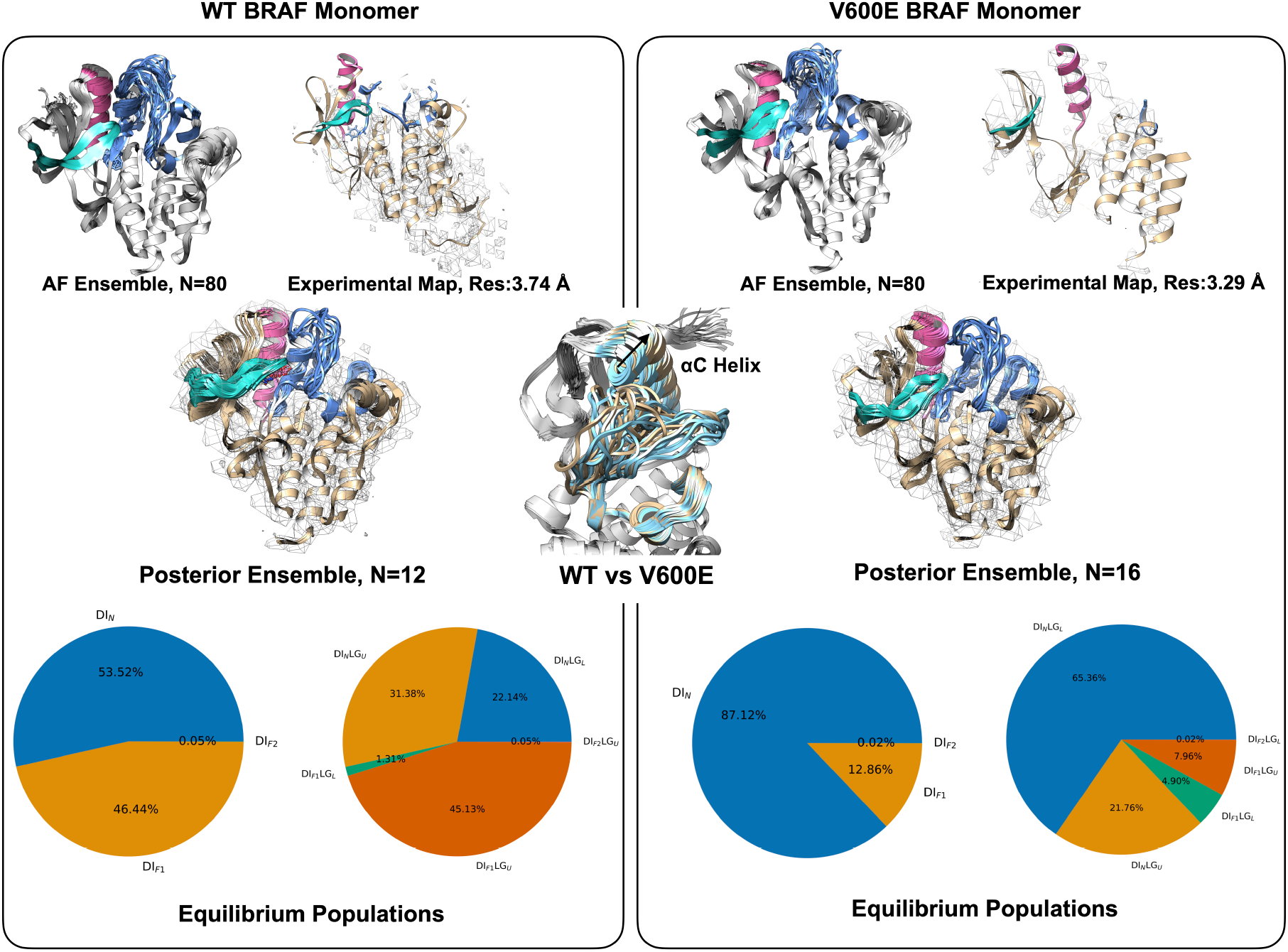
Cryo-EM structures of monomeric WT and V600E BRAF have been resolved in complex with the 14-3-3 scaffold. Using MSA subsampling, we generated conformational ensembles for both WT and V600E BRAF, capturing structural elements such as the activation loop (blue), αC-helix (magenta), and G-loop (sea green) that are absent from X-ray crystal structures fitted to cryo-EM maps. Bayesian reweighting of the AF-derived ensembles with reference maps produced posterior ensembles for WT and V600E that inherently represent the dynamic αC-helix and activation loop beyond the static snapshots of crystallography. Molecular simulations initiated from posterior ensembles and analyzed with MSMs revealed populations of multiple metastable states. MSM weighted equilibrium populations showed a clear shift from the DI_F1_ state in WT to the DI_N_ state in V600E, consistent with activation upon oncogenic mutation. This shift was reflected in the conformational dynamics of the αC-helix, which transitioned from αC-out (LG_U_) to (WT: 76.56%, V600E: 29.72%) αC-in (LG_L_) (WT: 23.45%, V600E: 70.28%) upon mutation.

**Figure 6.**
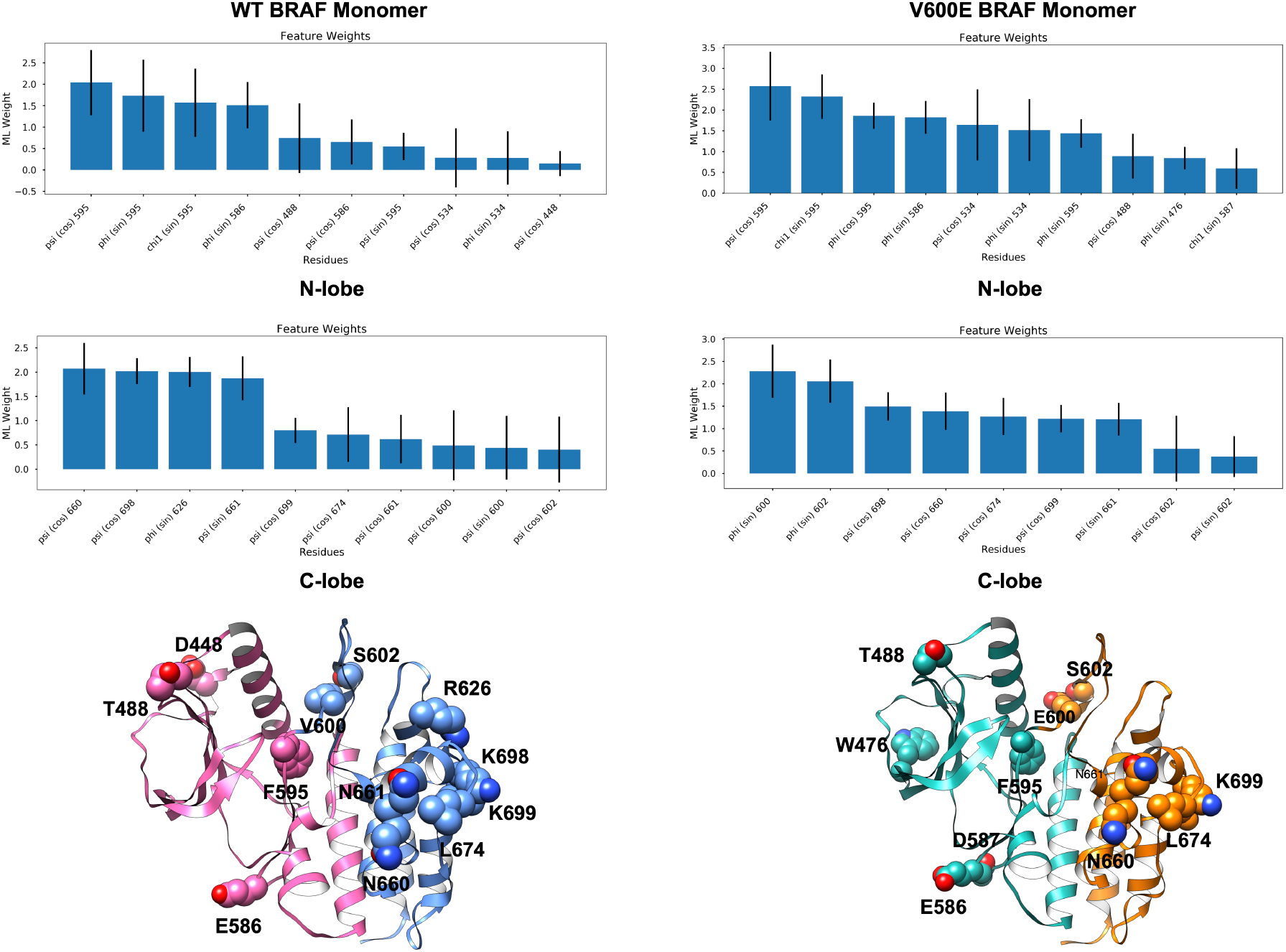
TAE trained on ϕ, ψ, and χ1 angles of the N-lobe (WT: magenta, V600E: sea green) and C-lobe (WT: blue, V600E: orange) regions of WT and V600E BRAF captured slowly varying structural features governing BRAF conformational dynamics. In both cases, the model identified DFG-Phe (F595) and V/E600 as key residues controlling these dynamics.

CryoPhold further enabled generation of a conformational ensemble for monomeric BRAF V600K, for which *Lavoie et al*. were unable to obtain a consensus structural representation (EMDB: EMD-43675, Res: 4.43 Å)^33^. Simulations initiated from the CryoPhold ensemble (total 12 structures in posterior ensemble totaling *12 * 3 * 200 ns* = 7.2 µs) revealed an *inactive* → *active* population shift similar to V600E, supporting the generalizability of mutation-induced population shifts in BRAF (Figure S3 in Supporting Information).

### Discussions

Our modular CryoPhold framework—merging AF2 generated conformational ensembles with cryo-EM density through Bayesian ensemble refinement, physics-based simulations, and machine learning, effectively captures populations of metastable states and conformational hotspots in both GlyT1 and BRAF. Multiple-sequence alignment (MSA) subsampling of AF2 or other generative AI,based structure prediction models has emerged as a powerful and accessible strategy for generating conformational ensembles of biomolecules. However, it remains an open question whether these models truly learn physical interactions, merely reproduce statistical patterns in the training set, or in some cases memorize the training data. Compounding this challenge, an averaged cryo-EM density map does not fully resolve conformational heterogeneity of biomolecular motions.

The Bayesian ensemble refinement approach employed here extracts diverse conformational states that best represent the average cryo-EM density, providing a principled way to incorporate experimental priors into AI-generated conformational ensembles. This addresses the long-standing limitation of fitting X-ray crystal structures into cryo-EM maps, where highly dynamic regions are often difficult to model accurately using X-ray crystallography. Our modular pipeline advances this further by embedding the CryoPhold refined ensemble with short molecular simulations and constructing Markov state models (MSMs), thereby linking static posterior ensembles to the underlying free-energy landscapes. The machine learning module overcomes the bottleneck of requiring prior structural knowledge to define low-dimensional coordinates^12^, while also identifying *slowly varying* structural hotspots that topologically connect functionally important metastable states.

A promising future direction is to use weights corresponding to each structure in the posterior ensemble to initiate a *weighted ensemble*^*37*^ workflow, where structures with higher posterior weights are prioritized as starting points for longer simulations. At the same time, lower-weighted states receive proportionally shorter simulations. This strategy ensures a robust scaling of computational resources enabling a sampling strategy that samples both dominant and rare but functionally important conformations.

CryoPhold has an intrinsic limitation. For larger molecular systems, the molecular dynamics simulations are computationally expensive and require significantly longer sampling times to capture key conformational transitions. In future, integrating population-aware diffusion generative models within the CryoPhold framework to alleviate this bottleneck.

To our knowledge, the CryoPhold workflow is the first of its kind to capture populations of metastable states in apo proteins and to quantify how disease-causing mutations modulate protein dynamics starting from cryo-EM–refined ensembles. Its modular design allows straightforward integration with emerging generative AI–based ensemble prediction models^38–40^ and collective variable discovery methods, enabling seamless unification of generative AI, cryo-EM, and enhanced sampling. Future efforts will enable integration of additional experimental priors, including single-molecule FRET^41^, DEER spectroscopy, and NMR^42^, using the BioEN framework in combination with enhanced molecular simulations^43^, thereby unifying statistical physics, biophysical experiments, and AI-based structure prediction in biomolecular modeling^44^.

### Methods

We developed CryoPhold as a modular pipeline. It begins by predicting a conformational ensemble with structural heterogeneity by modifying the MSA fed into generative AI–based structure prediction tools (“*Folding module*”). The ensemble is then refined by incorporating single-particle cryo-EM data using a Bayesian inference of ensembles (*BioEN*) approach (“*CryoModule*”), yielding a posterior ensemble. Independent structures from the posterior ensemble were used to seed unbiased molecular dynamics simulations (“*Simulation module*”). The resulting trajectory data are processed by a machine-learning (“*ML*”) module that uses high-dimensional data of protein dihedral angles and *Cα–Cα* distances as features and trains a time-lagged autoencoder to capture *slowly varying* structural motions and identify structural hotspots that govern conformational changes. ML-derived features together with manually identified features (aka collective variables) are then used to build a Markov state model (MSM), enabling prediction of the populations and transition times of metastable states in biomolecules that modulate function.

### Folding module

A structural ensemble for each sequence was generated using the ColabFold implementation of AF2 (link: https://colab.research.google.com/github/sokrypton/ColabFold/blob/main/AlphaFold2.ipynb#scrollTo=ADDuaolKmjGW), following the protocol outlined by *Meller and co-workers*^*45*^. An initial multiple sequence alignment (MSA) was constructed using the MMseqs2^46^ method integrated in ColabFold^47^. The MSA was then stochastically subsampled to include up to 8 cluster centers and 16 additional sequences (max_msa = 8:16). For ensemble generation, we used the “*greedy*” pairing strategy to match any taxonomically compatible subsets, ran 16 random seeds, and enabled model dropout. Leveraging both multiple seeds and dropout capitalizes on AF2’s model uncertainties, resulting 80 predicted structures per protein and capturing partial conformational heterogeneity.

### Filtering

The conformational ensemble generated by MSA subsampling was visually analyzed using UCSF Chimera^35^. For the apo GlyT1 structures, unrealistic conformations were filtered using pLDDT scoring. After testing several threshold values, a pLDDT cutoff of 50 was selected, which effectively removed unrealistic structures while preserving conformational diversity.

### Map generation and alignment

Structures generated from MSA subsampling were initially aligned to the cryo-EM map using the standalone Situs^48^ package. Focused reference maps corresponding to monomeric BRAF were generated in UCSF Chimera using the *vop zone* command, which selected the electron density within 5 Å of the monomeric kinase domain for both WT and V600E BRAF from BRAF-14-3-3 complexes.

### CryoModule

We generated a discrete structural ensemble 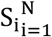 by MSA subsampling of AF2, i.e., drawing independent subsets of the multiple-sequence alignment and running separate predictions to obtain alternative conformations implied by sequence statistics. The experimental cryo-EM map *D*(*x*) on grid *x* ∈ Ω was clipped to nonnegative density and min–max normalized to [0,1]. A normalized threshold *τ* defined experimental support:

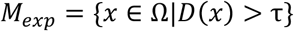

Each structure *S*_*i*_ was forward projected to a simulated density *M*_*i*_ (*x*) on the experimental grid, and two harmonization parameters were introduced: a global scale *s* > 0 and an isotropic Gaussian blur with standard deviation σ > 0, yielding:

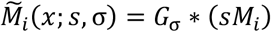

with * denoting convolution. A simulation-driven support *M*_*sim*_ was constructed from the ensemble (e.g., voxels exceeding several standard deviations across *M*_*i*_), and all voxelwise comparisons were restricted to the combined mask:

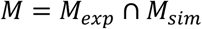

After masking and vectorization, we obtained *d* ∈ *R*^*V*^ from *D* and *m*_*i*_ ∈ *R*^*V*^ from 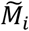, where *V* = |*M*|. Assuming additive Gaussian noise on masked voxels,

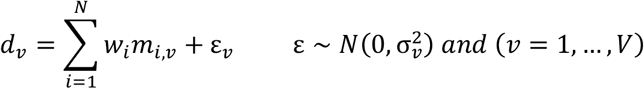

With σ_*ν*_ set on the normalized scale (e.g., 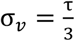), we inferred nonnegative normalized weights 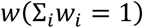 by minimizing the BioEN objective:

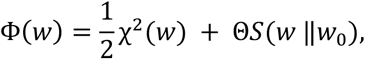

Where

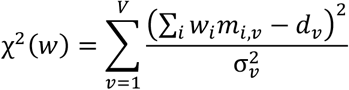

and

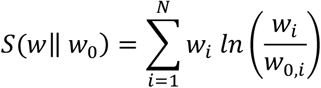

Here *w*_0_ is the uniform prior and Θ ≥ 0 controls regularization strength. Constraints were enforced via a softmax parameterization

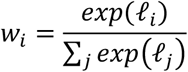

The nuisance parameters (*s*, σ) were fit to reconcile intensity and effective resolution, either as a prefit against the uniform average or within an alternating scheme minimizing

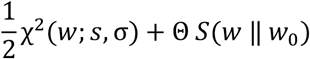

Regularization was chosen by tracing an L-curve over Θ and selecting the knee that balances fidelity and parsimony, quantified by the Shannon entropy:

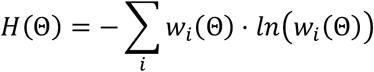

and the effective ensemble size

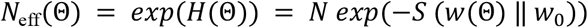

With the selected Θ* and weights *w*\*, we formed the posterior ensemble-average map

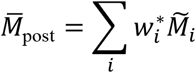

and the prior average:

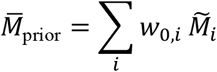

and evaluated agreement using real-space cross-correlation on *M* and Fourier Shell Correlation between *D* and 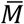, reporting the resolution at the first standard FSC crossing (e.g., 0.5).

### Simulation Module

Each structure from the *CryoPhold* posterior ensemble was prepared for molecular dynamics simulations using the *tleap* module in Amber2022^49,50^, following the protocols outlined by *Meller et al*.^*16*^ and *Vats et al*.^25^ Proteins were parameterized with the *AMBER ff14SB*^*51*^ force field. Counterions were added to each system, which was then solvated in a truncated-octahedron TIP3P^52^ water box, ensuring a minimum distance of 10 Å from any protein atom to the box boundary. Energy minimization proceeded in two phases: (i) solvent and ions were minimized while restraining the protein with a 100 kcal mol^−1^ Å^−2^ harmonic potential (200 steps steepest descent followed by 200 steps conjugate gradient); (ii) the entire system was minimized without restraints for 500 conjugate-gradient steps.

Following minimization in *Amber2022*, the Amber topologies were converted to GROMACS format using Acpype^53^. Each system was then gradually heated from 0 K to 300 K over 500 ps in the NVT ensemble, applying harmonic restraints (500 kJ mol^−1^ nm^−2^) on backbone heavy atoms. Restraints were subsequently removed, and the systems were equilibrated for 200 ps in the NPT ensemble at 300 K, with pressure maintained at 1 bar by a Parrinello–Rahman^54^ barostat and temperature controlled by a v-rescale thermostat. Production simulations were conducted in the NPT ensemble at 300 K and 1 bar using the leapfrog integrator with a *2 fs* time step. A 1.0 nm cutoff was used for nonbonded interactions, while long-range electrostatics were treated with particle-mesh Ewald (PME) using a 0.16 nm grid spacing. The *LINCS* algorithm^55^ constrained covalent bonds to hydrogen atoms. All heating, equilibration, and production runs were carried out using GROMACS 2022. Trajectories were saved every 10 ps, and the lengths of production runs are highlighted in the “*Results*” section.

### Featurization

#### Dihedral angles

Dihedral angles generated from CryoPhold-augmented molecular simulations were transformed using sine and cosine functions to resolve the inherent periodicity of angular variables. “*dihedral*.*py*” function of CryoPhold was used to extract user-defined dihedral angles from molecular dynamics dataset.

#### Distances

To convert C_*α*_ – C_*α*_ distances from MD trajectories into smooth and differentiable features suitable for state definition, we employed a contact switching function. For each *Cα-Cα* distance *r*, the transformation is defined as:

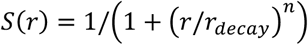

where *r*_*decay*_ is a decay distance parameter and *n* is an exponent controlling the steepness of the transition. In this formulation, short distances (*r* ≪ *r*_*decay*_) yield values close to 1, indicating a “*formed contact*”, while long distances (*r* ≫ *r*_*decay*_) yield values close to 0, indicating “*no contact*”. Unlike sharp cutoff functions, this continuous mapping reduces sensitivity to noise and provides a probabilistic measure of contact strength.

For this study, we set the decay distance to *r*_*decay*_ = 0.90 *nm* with *n* = 6. Distance trajectories were obtained from precomputed molecular simulations and transformed using this switching function, resulting in time series of contact probabilities *S*(*r*) bounded between 0 and 1.

To discretize these contact probabilities into interpretable states, we applied two thresholds in the switching-function domain:

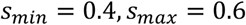

Contacts with *S*(*r*) ≥ *s*_*max*_ were classified as formed, while those with *S*(*r*) ≤ *s*_*min*_ were classified as broken. Values within the transition window (*s*_*min*_ < *S*(*r*) < *s*_*max*_) were treated as intermediate states, reflecting dynamic fluctuations where contact formation is ambiguous. This thresholding strategy balances robustness against random noise with sensitivity to meaningful conformational transitions.

Together, the switching-function transformation and thresholding procedure provide a smooth, interpretable framework for representing contact dynamics, enabling downstream dimensionality reduction.

#### ML Module

ML module consists of two algorithms which reduces the high-dimensional data generated by molecular dynamics simulations to generate a low dimensional representation and also allows identification of structural hotspots governing protein dynamics.

#### Slow Feature Analysis (SFA)

Slow Feature Analysis (SFA)^25^ in context of biomolecular simulation was introduced by *Vats et al*. is a dimensionality reduction technique designed for high-dimensional temporal data. Its primary objective is to transform the *J*-dimensional input signal, *c*(*t*), into output signals *y*_*k*_(*t*) = *g*_*k*_(*c*(*t*)) using a set of nonlinear functions gk(c). These output signals are optimized to minimize

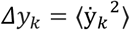

where 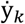 represents the temporal derivative of *y*_*k*_(*t*), and ⟨·⟩ denotes temporal averaging. By minimizing Δ*y*_*k*_, SFA extracts features that vary slowly over time. In this study, we demonstrate how SFA can be applied to identify *slowly fluctuating* distance pairs which captures the conformational *hotspots* from CryoPhold augmented molecular simulation.

#### Time-lagged Autoencoder

The time-lagged autoencoder (TAE)^22^ was designed to learn compact representations of time-series data while explicitly leveraging temporal structure. This approach is particularly powerful in scenarios such as molecular dynamics (MD) simulations, where the goal is to capture slowly varying conformational dynamics of biomolecules from high-dimensional data.

The TAE was introduced previously by *Bhakat et al*.^*23*^ and comprises a) Encoder: Transforms the current system state *x*_*t*,_ into a lower-dimensional representation *z*_*t*._ b) Decoder: Reconstructs the time-lagged version of the system, *x*_*t*+*τ*_, from the latent representation *z*_*t*_. By pairing *x*_*t*_ and *x*_*t*+*τ*_, the model naturally emphasizes slowly varying dynamics, since the most prominent changes occurring over lag time τ are embedded in the latent space.

Latent Layer Representation: Let *x*_*t*_ ∈ ℝ^*d*^ be the high-dimensional molecular state at time t. In the encoder, the latent layer is often formulated as:

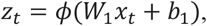

where:

- *W*_1_ ∈ ℝ^(*m*×*d*)^ is the encoder’s weight matrix.
- *b*_1_ ∈ ℝ^*m*^ is the bias vector.
- *ϕ*(·) is a nonlinear activation function.
- *z*_*t*_ ∈ ℝ^*m*^ is the low-dimensional latent representation.

The decoder then reconstructs *x*_*t*+*τ*_ from *z*_*t*_, typically using:

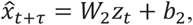

Where *W*_*2*_ ∈ ℝ^(*d* × *m*)^ and *b*_*2*_ ∈ ℝ^*d*^. The training objective is to minimize the reconstruction error:

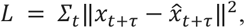

which encourages *z*_*t*_ to preserve information about the slow degrees of freedom predictive of future conformations.

The TAE’s learned latent space provides a compressed, nonlinear representation of input features that captures slowly varying conformational dynamics in molecular systems. The framework accepts sin–cos– transformed dihedral angles and Cα–Cα distances as inputs, ensuring that all relevant motions are incorporated into the dimensionality reduction. Our machine learning approach applies a dynamic feature-selection strategy using a learnable masked kernel with *ReLU* activation, retaining only features with non-zero weights. This provides two advantages: the retained features are more interpretable, and they can be directly used as collective variables (CVs) in enhanced sampling simulations.

To further structure the latent space, two extensions were introduced: (a) Gaussian noise was added to the encoder output prior to decoding, encouraging separation of long-lived states; and (b) the training objective was expanded with the VAMP score^56^ computed in latent space, which ranks states according to the slowest dynamical processes.

During training, the “*CVEstimator*” module balances three terms—reconstruction loss, a penalty on large masked-kernel weights, and the *VAMP* loss—each weighted appropriately to optimize both interpretability and dynamical fidelity. This integrated design yields an end-to-end pipeline where feature selection, sparse representation learning, and dynamical analysis collectively provide a robust and physically meaningful representation of the input data. In this study, TAE was used to identify slowly varying dihedral angles from the high-dimensional temporal data generated by MD simulations seeded from CryoPhold generated posterior ensemble.

#### Markov state modelling

MSM was performed on low-dimensional representations extracted from molecular simulations using PyEMMA 2.5.7^57^.

## Supporting Information

Free energy surfaces, populations of conformational states across different systems (WT, V600E and V600K BRAF) are available in the Supporting Information.

## Software Availability

CryoPhold can be accessed here: https://github.com/strauchlab/cryoPhold

## Author Contributions

SB: Conceived the idea, performed molecular simulations, dimensionality reduction, and wrote the initial draft. SV: Developed the cryomodule and slow feature analysis framework. AM: Developed the time-lagged autoencoder framework. EMS: Supervision, refining the idea, computing resources and co-writing the manuscript.

