## Supplementary material for "CryoPhold: CryoEM meets AlphaFold and molecular simulation to reveal protein dynamics": supp. info


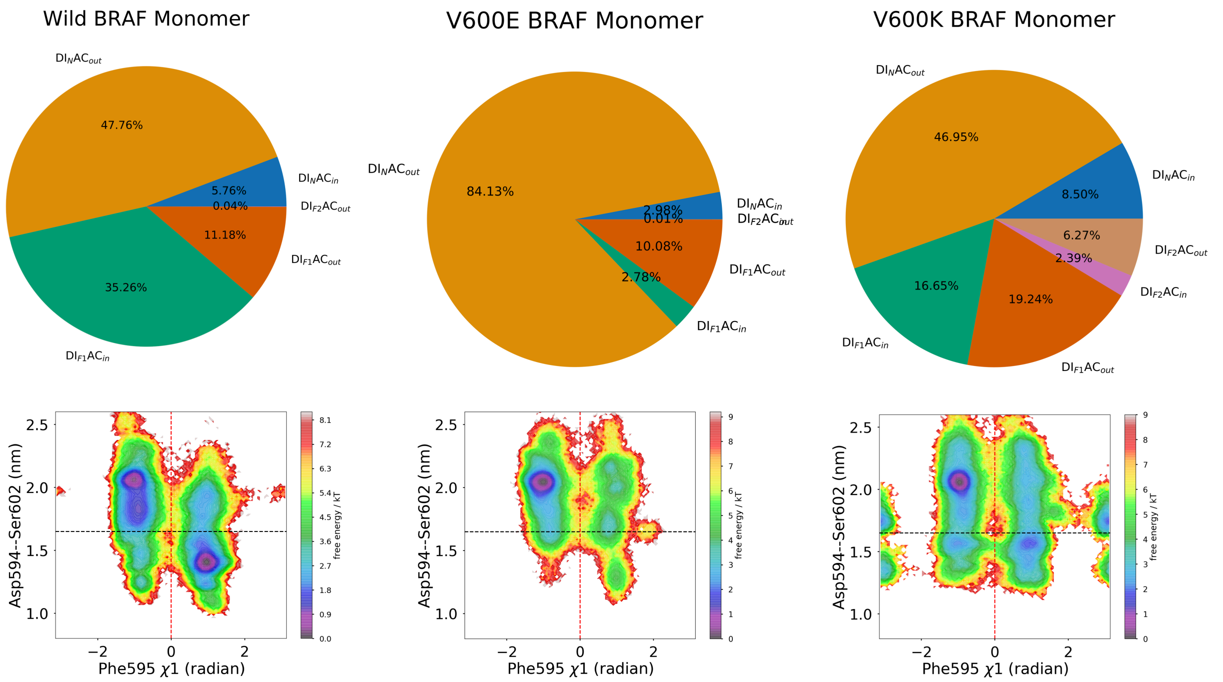


**Figure S1**. Upper panel: MSM-weighted populations of DFG-Phe and activation loop states show how the oncogenic V600E and V600K mutations shift the activation loop from folded ($AC_{in}$) to extended ($AC_{out}$) conformations. Lower panel: Population-weighted free energy surface projected along the DFG-Phe dihedral angle and the Cα–Cα distance between Asp594 and Ser602 highlights this population shift, governed by DFG-Phe flipping and activation loop dynamics.

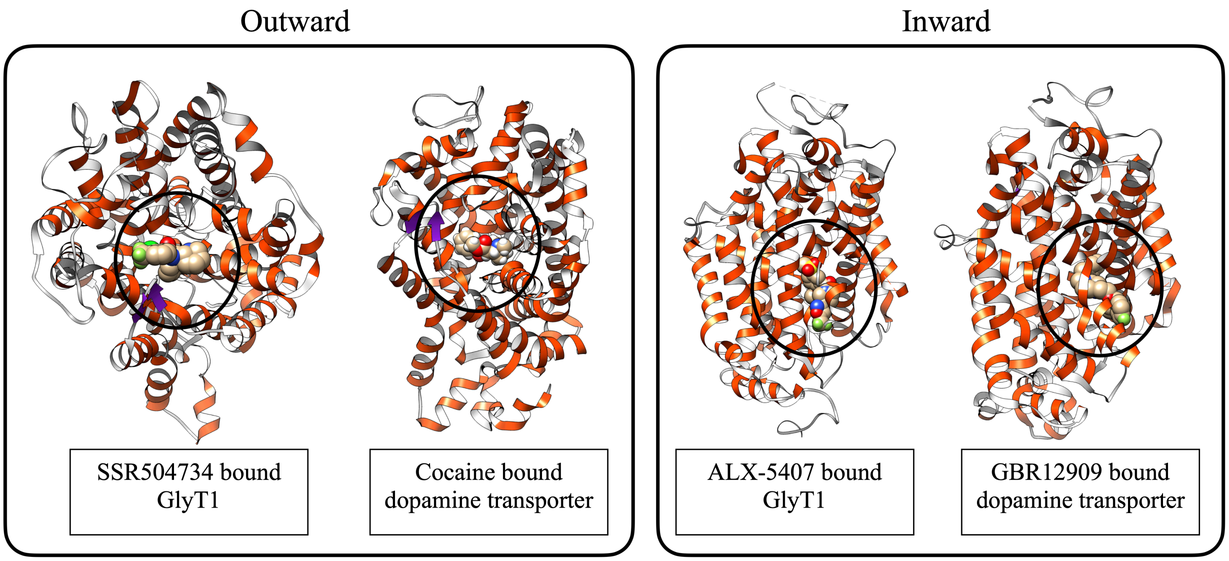


**Figure S2**. Top view of holo human transporters, GlyT1 and dopamine transporter, bound to small molecules highlights the exposed binding pocket, which stabilizes the outward conformation. The side-view of GlyT1 and dopamine transporter bound to small molecules ALX-5407 and GBR12909 shows an allosteric binding site buried within transmembrane helices.

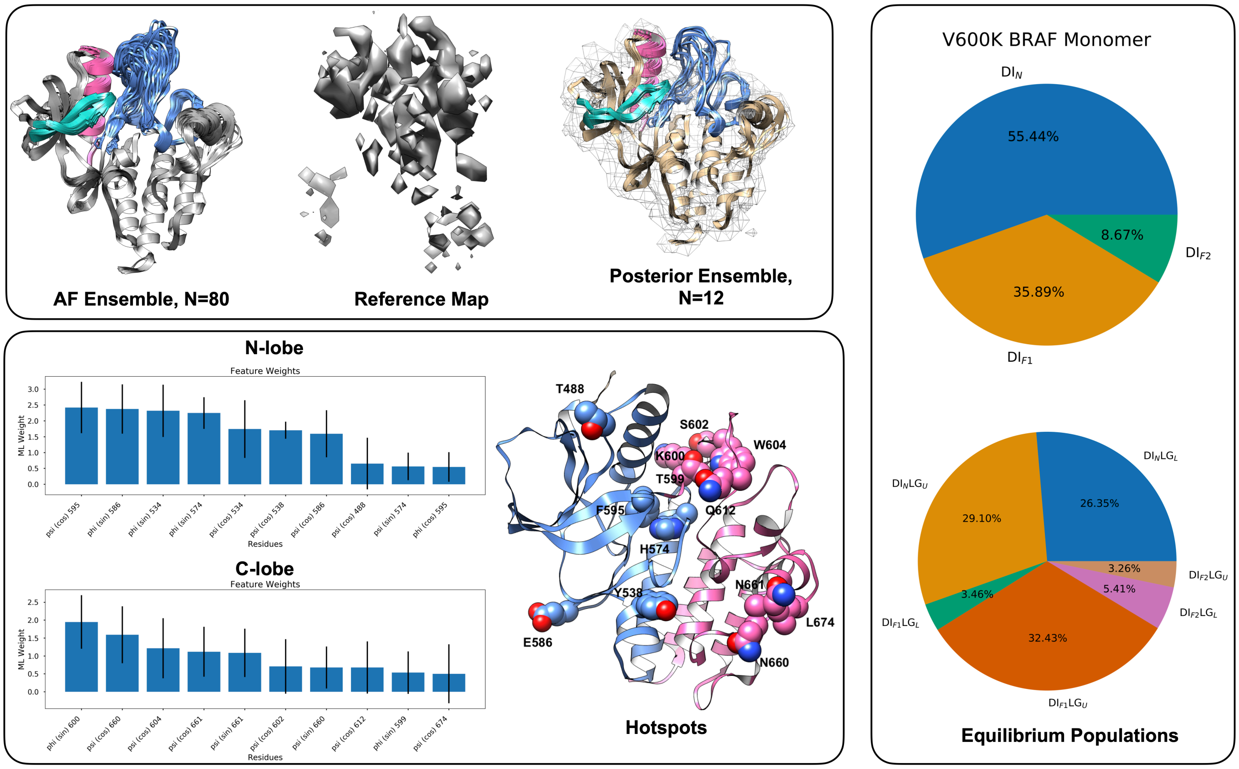


**Figure S3**. MSA subsampling generated a conformational ensemble of monomeric BRAF V600K, highlighting key structural regions including the activation loop (blue), αC-helix (magenta), and G-loop (sea green). Notably, *Lavoie*

*et al.* were unable to obtain an X-ray crystal structure corresponding to the reference cryo-EM map for the V600K mutant (EMDB: EMD-43675, Res: 4.43 Å). Molecular simulations initiated from this posterior ensemble revealed distinct metastable populations, with V600K showing a reduced fraction of the $DI_{F1}$ state compared to wild type BRAF (*V600K: 35.89%; WT: 46.44%*), thereby capturing the mutation-induced population shift. Machine learning analysis of the simulation trajectories identified key structural determinants ("Hotspots") of protein dynamics within the N-lobe (blue) and C-lobe (magenta). Importantly, the model consistently recovered DFG-Phe (F595) and K600 as critical residues governing the slow, timescale-defining motions in V600K BRAF.
